# Repetitive transcranial magnetic stimulation increases phase-amplitude coupling

**DOI:** 10.1101/134239

**Authors:** Chie Nakatani, Caitlin Mullin, Johan Wagemans, Cees van Leeuwen

## Abstract

A prominent feature of brain activity with relevance to cognitive processes is Phase-Amplitude Coupling (PAC) between slow and fast oscillatory signals. A newly developed neural mass model of cross-frequency coupling [1] predicts, counter-intuitively, that PAC shows sustained increases after repetitive Transcranial Magnetic Stimulation (rTMS). This is because rTMS leads to simultaneous neuronal firing in distinct regions, thereby enhancing the connectivity that, according to the model, is needed for PAC to be increased. We tested this prediction in healthy human volunteers. Two seconds of 10Hz rTMS were applied to the intraparietal sulcus, temporal-parietal junction, and lateral occipital complex. PAC in the subsequent electro-encephalogram was analyzed for two band pairs, theta-gamma and alpha-gamma, and compared to a sham condition. For all stimulation loci, PAC was higher in both band pairs after rTMS than in the sham condition. These results were found to be conform the model prediction. The perspective for using rTMS to modulate cross-band coupling is discussed.

## Introduction

Field signals from the brain, such as, EEG, MEG, ECoG, and LFP, all comprise multiple frequency bands, including, from slow to fast: delta (1-4Hz), theta (4-8Hz), alpha (8-13Hz), beta (13-30Hz), and gamma (> 30Hz) bands, as well as frequencies beyond these conventional bands [2]. The frequency composition of the signals as expressed by their power bands not only differs across brain regions [3] but also changes dynamically, for instance in switches between resting vs. task-related states. The diversity and dynamicity are a product of interactions between various types of neurons in cortical and subcortical regions [4-6]. The morphological, electric, and synaptic diversity of the neurons notwithstanding, we may distinguish two principled ways with which the multiple frequency components are generated [7]. One is through the interactions of excitatory and inhibitory neuron populations [8-10]. The resulting oscillation frequency is largely determined by the response properties of inhibitory neurons, in particular their latencies. The variety of their response properties [11], thus, generates a wide frequency band of oscillations. Another mechanism responsible for oscillation is self-feedback of inhibitory neurons [12]. This mechanism is observed in inhibitory interneurons through which fast oscillations, namely in the gamma band frequency range, are generated [13, 14]. In a small volume of cortical tissue, both types of oscillators are connected [15, 16]. They interact non-linearly, resulting in cross-modulation components that are a product of the two bands [1].

Such modulation components are hard to detect in the power spectrum; in the frequency domain, they are often buried in 1/f noise. Therefore, in order to detect/assess cross-band modulation, measures based on time-frequency domain signal are used, e.g., phase-amplitude coupling (PAC) between bands of interest. Of possible band pairs, modulation from a slow band, such as theta and alpha, to gamma band activity has been investigated most intensively (see [7] for a review). Empirical studies have provided enough anatomical and physiological details to sketch the neural mechanism for the modulation; Spike timing of neocortical pyramidal neurons is primarily controlled by inhibitory interneurons, especially, by those with fast-spiking ability [16]. These interneurons synchronize their activity in the gamma band; thus, spiking of multiple pyramidal neurons is also synchronized in that band. The interneuron gamma activity is modulated in amplitude by the phase of slow band, e.g., theta activity [17-20]. Since the interneurons make synaptic connection to pyramidal neurons, spiking behavior of pyramidal neurons also shows this modulation in their gamma synchrony. As a result, field signals from a small piece of cortex contain modulated gamma band oscillation, which could be considered as an index of population spiking activity, e.g., [21].

PAC between the slow and gamma bands is observed, not only within a local region, but also across brain regions [22, 23]. Such ensemble activity over brain regions has been considered crucial for high-level mental activity [24-26]. A simulation study showed that cross-regional modulation could emerge in a network of oscillators which are capable of generating local phase-amplitude modulation to the gamma activity [27]. Taken together, PAC between the slow and gamma activities offers a possible mechanism for coordinating spike-timing across regions to implement cognitive functions [28].

A good number of studies have reported correlation between the PAC and psychological variables. For instance, the PAC was higher with than without a cognitive task [29]. PAC strength changed with attention [18] and correlated with task performance[30-32]. These studies suggest that PAC is involved in high-level cognitive functions and behavior. Moreover, the coupling could implement long-term effects, such as practice effect on task performance[32]. PAC strength, therefore, could substantiate both transient and persistent effects in cognition and behavior.

We may wonder whether PAC strength is affected by external manipulation, such as application of transcranial magnetic stimulation (TMS). To our knowledge, no studies have addressed this issue. This may, in part, stem from the difficulty of what effect to expect. Although the TMS pulses are focused on a small patch of cortex, it still contains tens of thousands of neurons. Given that neural oscillations (and PAC) emerge as a result of interaction among these neurons, it is very hard, if not impossible, to predict what effect TMS would have, unless a model could describe the effect on PAC at population-level.

Recently, we proposed a neural mass model of PAC[1, 27]. The core module of the model consists of four units modeled after the population dynamics of pyramidal neurons, slow inhibitory, fast inhibitory, and excitatory interneurons in cortex. The four-populations module represents a cortical patch between a few millimeters and a centimeter in diameter, in which oscillations in various frequencies can be observed [33], among which PAC may occur. In particular, the model predicts, counter to what one would intuitively expect, that PAC does not decrease, but *increases* when a typical TMS is applied.

The interaction between pyramidal and slow inhibitory interneurons in the model generates the slow main activity. The fast-inhibitory interneurons have dynamic self-feedback, which generates gamma band oscillation. Input noise level determines the state of the fast frequency; as the level increases moderately, the fast oscillation emerges, and is stabilized. With further increase, the fast oscillations would eventually disappear due to saturation. The presence of fast oscillations is a necessary condition for PAC to occur. PAC arises in the model through couplings between the pyramidal/slow inhibitory interneuron and fast inhibitory interneuron populations. Thus, gamma oscillation is amplitude-modulated by the slow activity. Specifically, according to the model, the input noise level and the connectivity jointly determine the level of amplitude-modulation of gamma oscillation is by the slow activity.

The four-populations model predicts that PAC does not decrease, but *increases* when typical TMS is applied. Typical strength of TMS pulse is near firing threshold (e.g., 110% of motor threshold) to keep the stimulation effective but low-dose for participants. In such case, the input noise level would increase moderately, which, according to the model, would result in increase in gamma band power. However, this effect is only transient; the PAC level goes back to baseline as soon as the stimulation is turned off. In contrast, the connectivity parameter of the model is responsible for long-term effects on PAC strength. Our current focus is on long-term modulation on PAC intensity which corresponds to behavioral practice effect [32]. Thus, the connectivity was considered as the primary target in the current study.

TMS forces neuronal populations to fire simultaneously. When the stimulation is repeated (rTMS), Hebb’s law prescribes increase in connectivity strength among the neurons. This results in an increase in PAC strength. Thus, the model predicts a sustained increase in PAC after rTMS. In the post-rTMS period, the effect of input noise level should be negligible. Therefore, no increase in gamma power is expected. To test these predictions, we applied rTMS and analyzed PAC of post stimulation periods (10s/period).

## Methods

### Participants

Fifteen young adults (average 22.33 years old, 8 female) participated in the experiment. The participants had no history of major illness, and normal or corrected-to-normal vision. All of them were right-handed. They gave a written informed consent to the current study which was approved by the ethics committee for medical scientific research at the KU Leuven.

### Anatomical MRI acquisition

A T1 weighted-anatomical image was obtained for each participant. The image was used to determine TMS target locations and to localize EEG sources. The image was acquired using a 3T Philips Ingenia CX scanner at the Department of Radiology of the KU Leuven with a 32 channel head coil. One hundred eighty two contiguous coronal slices were obtained using 3D-turbo field echo (3DTFE) sequence (time repetition (TR) = 9.6 ms, time echo (TE) = 4.6 ms, field of view (FoV) = 250 x 250 x 218 mm, time inversion =863.4 ms, slice thickness= 1.2 mm, and in-plane resolution = 0.98 x 0.98 mm).

### TMS target selection

Three brain regions were chosen as rTMS targets: intraparietal sulcus (IPS), temporal-parietal junction (TPJ), and lateral occipital complex (LOC). These regions are relevant for visual target reportability and perception [34-37]. Another key region, dorsolateral prefrontal cortex was not included in the current study because a pilot study showed that stimulation to the frontal region induced large and wide-spread muscular and ocular artefacts. The anatomical MRI was processed with BrainVoyager QX v2.8 (Brain Innovation, Maastricht, the Netherlands). The image was re-sampled to 1x1x1 mm voxels, then further transformed to the Talairach coordinate system, in which stimulation target locations were determined. The three targets were on the right hemisphere. The coordinates were [X = 28, Y = −51, Z = 50] for IPS, [X = 42, Y = −62, Z = 17] for TPJ, and [X = 40, Y = 75, Z=13] for LOC (Figure 1). The coordinates were decided based on previous studies ([38]for IPS, [39]for TPJ, and [35]for LOC). For participants in whom the coordinates do not correspond to gray matter, the target location was manually adjusted to the gray matter closest to the coordinates. Target markers were placed in the image. The image was then back-transformed to real 3D space. The MRI file with target markers was used in the BrainSight Neuronavigation system (Rogue Research, Montreal, Canada) to guide the TMS coil to the stimulation sites.

**Figure 1.**
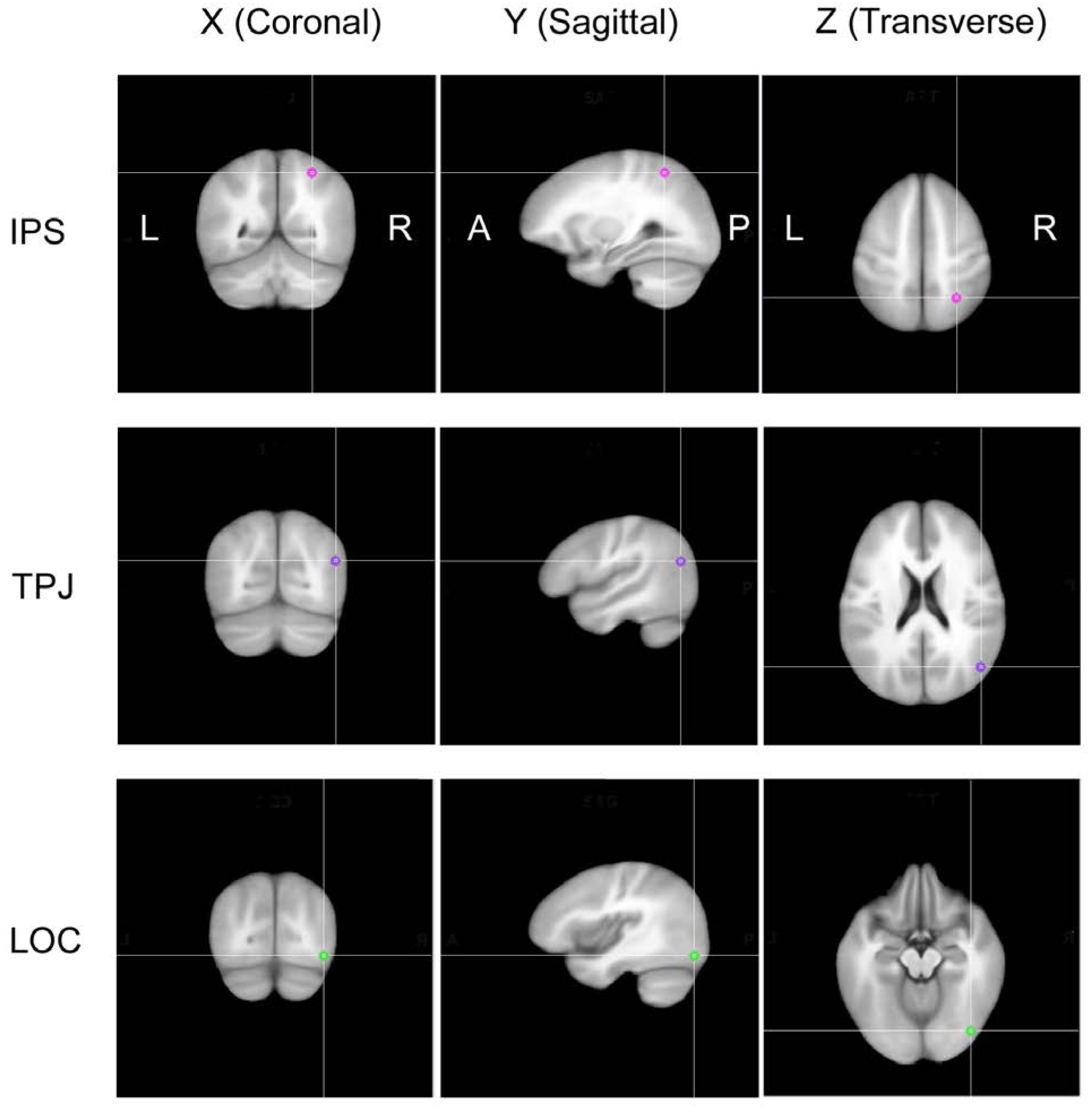
Stimulation loci. Three stimulation loci in right intraparietal sulcus (IPS), temporal parietal junction (TPJ), and lateral occipital complex (LOC) were shown on ICBM brain (average over T1-weighted images of 452 subjects) in Talairach coordinate.

## TMS equipment and stimulation parameters

TMS was applied using the Rapid^2^ system (Magstim, Whitland, UK). A letter-eight coil was used to administer the magnetic pulses. Pulse intensity was decided based on motor threshold session to left motor cortex to five participants (two of them are participants in the current study, while three participated in a pilot). For a TMS pulse to be super-threshold, 110% of the motor threshold value was used for the two of participants who also underwent the motor threshold session. For the other participants, the median of the five super threshold values, which was 65% of default output power of the stimulator, was used.

An rTMS pulse train consisted of 20 TMS pulses in 10 Hz (i.e., 2 s), Stimulation number and frequency were chosen based on a literature review [40]. After the stimulation, 10 s no-stimulation period was inserted. The 10 s post-rTMS period was used for data analyses (Figure 2). The rTMS trial was repeated 20 times to each stimulation site. A sham condition used the same rTMS and 10 s break schedule with the coil rotated 90 degrees. Thus, in the sham condition, magnetic stimulation was not delivered to the brain, but the clicking sounds and electromagnetic noise due to the TMS were comparable to those in the rTMS condition. The sham condition was recorded prior to the rTMS condition at each stimulation location.

**Figure 2.**
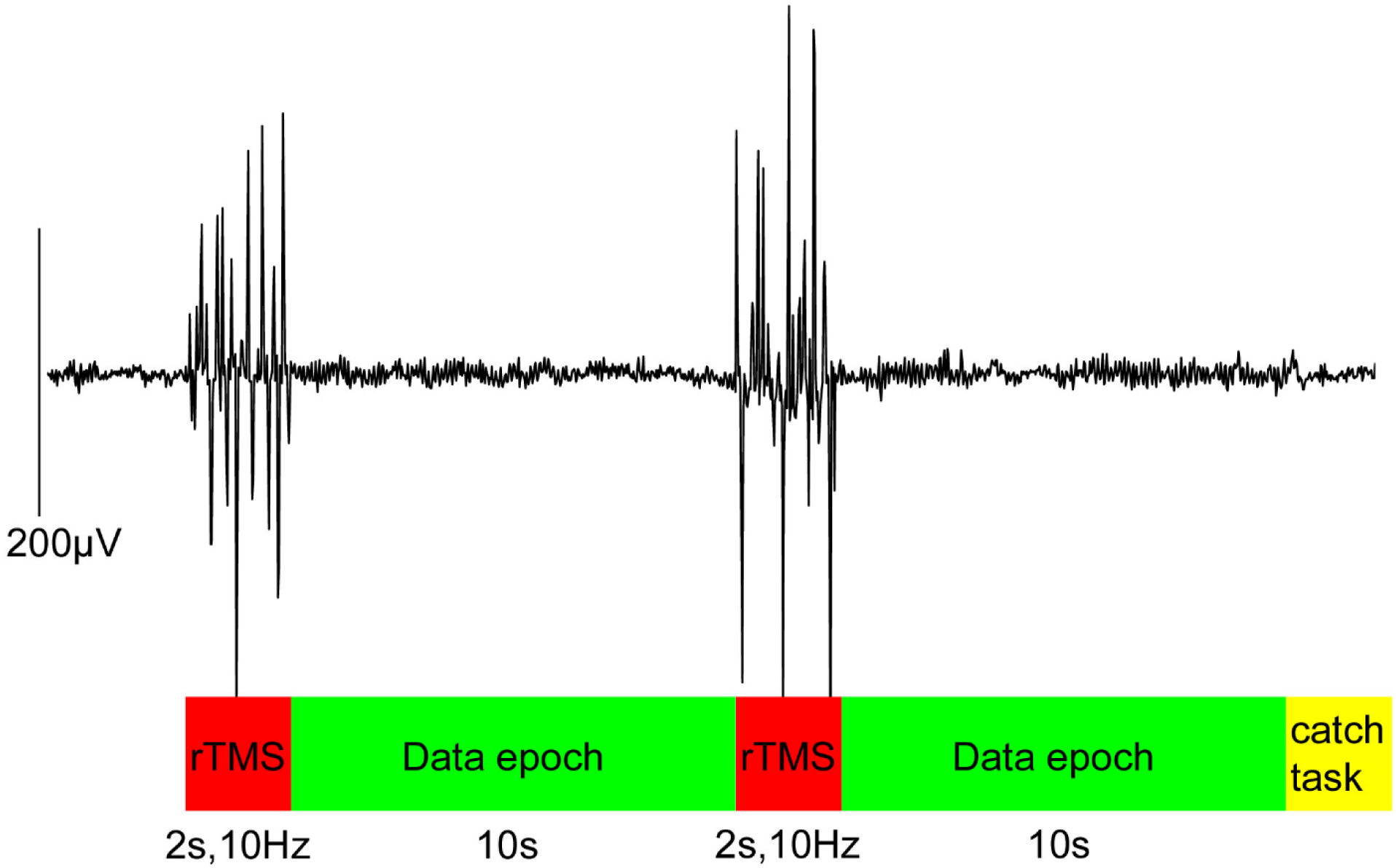
rTMS and data epoch. Twenty pulses of TMS was applied in 10 Hz. Ten seconds from the offset of rTMS was used as data epoch. Display change for catch task occurred every 5 iterations of rTMS and data epoch. Participants pressed a button to report the change. The catch task period was not included to data analysis.

### EEG equipment and recording settings

EEG was recorded using a TMS compatible system (Waveguard Duke64 cap and Patientguard EEG recording system, ANT Neuro, Enschede, The Netherlands). Sixty five electrodes were placed in an approximately equidistant hexagonal/pentagonal layout, a configuration suitable for source modeling. The number includes two channels for vertical and horizontal EOGs. The right earlobe was the recording ground. Recording range was 0.5 – 200 Hz. Sampling rate was 2048 Hz.

### Task and experiment control system

To control participants’ vigilance level and eye fixation, a visual detection task was used. The participants fixated a white crosshair at the center of the screen (Dell 1070FP, refresh rate was 60Hz). When the fixation changed the color to blue, participants were asked to press the ‘5’ button on a small ten-key pad. Experimental task presentation, response button press, TMS trigger, and EEG marker signals were controlled by Presentation v.14.8.12.30.10 on a Windows XP PC (Dell XP Professional).

### Procedure

The anatomical MRI and the rTMS & EEG sessions were held on two different days. The MRI session was held one week or more prior to the rTMS & EEG session. In the rTMS & EEG session, participants wore the EEG electrode cap. Conductive gel was injected to each electrode. Electrode impedance was kept below 5 kOhm. Markers for head tracking were attached on the electrode cap. The participant was then seated in front of the computer display in which task-related stimuli were presented. The participant used a chin rest to hold the head as stable as possible during the experiment. Distance between the chin rest and the center of display was constant within a participant, but was different between participants (67 - 75 cm) depending on individual body size.

Once the participant’s head position was secured, co-registration of head fiducial points in the real space and a head image reconstructed from MRI was performed. Spatial location of the nasion, tip of nose, right and left preauricular points were sampled by the pointing devise as the fiducials. For some participants whose nose tip was not available in the MRI image, corners of eyes were used as alternative points. The co-registration was repeated when necessary, e.g., when the participant’s head moved significantly. To compensate for small head movements, TMS coil orientation was monitored during stimulation via the BrainSight Neuronavigation system. Coil orientation was manually adjusted to keep the tracking error within a 2.5 mm radius from the target location. Segments with significant tracking error were excluded from analysis.

The rTMS and sham conditions were part of a basic data acquisition session, which also included resting, visual evoked, and single-pulse TMS conditions. Total duration of the session was 3-4h, including breaks between the conditions. The single-pulse TMS was applied prior to the rTMS condition at each of the three locations. One hundred single pulses were administered with an SOA between 2000 and 2400 ms. The pulse intensity was the same as that of rTMS. Our pilot study showed that the evoked response of single-pulse TMS came back to baseline by 500ms. A break (∼20 min) before an rTMS session was, therefore, sufficient to exclude the effect. In principle, the single-pulse TMS could be considered as low-frequency rTMS (∼0.5 Hz). Therefore, it could have the same effect of increasing connectivity as predicted for the rTMS condition, an effect that could extend beyond the 20-minute break.

## Results

### Results of catch task

During the rTMS & EEG sessions, the participants reported the color change of the fixation cross with 100% accuracy. Participants, therefore, were wakeful and alert during the sessions. Segments of EEG recordings between the display change and 500ms after the button press were excluded from the following EEG analyses.

### EEG Preprocessing

To reduce noise from TMS pulses, an EEG segment between -20 and +70 ms from pulse onset was cut, and the gap was linearly interpolated using a Python script. Residual TMS and other noises, such as EOG, EMG and mains noise, were reduced further using a commercial EEG signal analysis package (BrainVisionAnalyzer ver. 2.1, Brain Products, Germany). The data was down-sampled from 2048Hz to 512Hz. The signal was low cut at 1.0Hz and high cut at 45.0 Hz using an IIR/no phase-shift filter. Independent component analysis (InfoMax algorithm) was then applied to identify EOG, EMG, mains, and residual of TMS noise components. The noise components were visually identified and excluded from signal reconstruction. The cleaned data was cut to 10 s segments from the end of rTMS. The segments were checked once more using the semiautomatic artifact rejection routine of the analysis package. Two participants had a few noisy channels (two for S4, three for S5; mostly; mostly frontal and central electrodes), which were replaced by the average of signals from neighboring channels. At the end of the preprocessing procedures, 11.6 % of epoch segments, including all epochs of one participant, were discarded. The main data analysis was conducted on the clean epoch segments of the 14 remaining participants: on average over conditions and stimulus locations, 16.75 epochs (167.5 s) were used.

### Source estimation

From the cleaned EEG sensor signals, cortical activity was estimated using distributed source estimation methods implemented in Brainstorm [41, 42]. The estimation requires cortical and head surface information from each participant. The surfaces were reconstructed from T1-weighted image using Freesurfer v5.3.0 (http://surfer.nmr.mgh.harvard.edu/). The structural information was co-registered to the EEG data using Brainstorm functions. Fiducial points were the nasion, bilateral preauricular points, and an arbitrary inter-hemispheric point. Moreover, eight electrodes were used to adjust the electrode location on the scalp manually: the electrodes located above eyes (1L and 1R; electrode labels are *not* according to the international 10-10, but unique to the cap), in front of ears (2LD and 2RD), behind ears (3LD and 3RD), and around the vertex (4Z and 5Z). A head model was computed based on the boundary element method (BEM) using OpenMEEG [43, 44] incorporated in Brainstorm. Three boundary elements, scalp, inner skull, and outer skull had 1922 vertices per boundary.

Prior to the estimation, EEG signals were re-referenced to the average reference. The period between 0.5 and 9.5s was used in each epoch to avoid any effect of segment edges and residual TMS evoked responses. The noise covariance matrix, which shows the correlations between sensor signals at baseline, was computed from one of the sham epochs. Thus, the epoch was excluded from source estimation. The minimum-norm estimation method was used to estimate distributed source activity. Possible source locations were restricted to the cortical surface. The surface was estimated by the triangular mesh computed from the anatomical MRI image. A total of 15002 points on the vertices of the mesh were used. At each point, three orthogonal dipoles were placed to model local activity. Source activity was estimated for each single epoch.

### Estimated signals of the TMS regions

TMS stimulation location could jitter due to head motion and tracking lag/error of the TMS coil. Also, error in source localization should be considered. Based on these considerations, we chose 50 sources (dipole triplets) at around each TMS location for the analytical signals. In each participant, a dipole triplet closest to the TMS target coordinate was chosen as the seed, and adjacent sources were added up to 50 triplets to provide a region of interest (ROI). The ROI covers approximately 10cm^2^ of cortex. There was no overlap among the three ROIs within a participant. Using source analysis functions offered by Brainstorm, the norm of each triplet was computed and averaged over the 50 sources in each post rTMS epoch. Single epoch ROI signals were used for the power spectrum and phase-amplitude coupling analyses.

### Power spectrum analysis

Power spectrum was computed using the Welch method as implemented in Brainstorm. The window length was 1s, and window overlap ratio was 50%. The power spectrum was computed from single epochs. The single trial power spectra were averaged over epochs for each stimulation condition within participant; then a grand average over the participants and standard error (SE) were computed. Figure 4 shows the grand average power and SE of the rTMS and sham conditions for each stimulation locus. We tested the difference between the rTMS and sham conditions in five frequency bands; 2-4Hz as delta, 5 - 7 Hz as theta, 8 - 12 Hz as alpha, 15 – 29 Hz as beta, and 30 - 45 Hz as gamma bands. Within each band, power was log transformed and averaged in each individual. The individual band average was submitted to statistical tests. Variance over participants was higher for the lower bands. Due to the unequal variance across the bands, the effect of rTMS was tested in each band. R (ver, 3.3.1, R Core Team, 2016) was used for all statistical tests. A linear model, Stimulation (rTMS vs. sham) by Loci (IPS, TPJ, and LOC) was fit in each band power data. In delta, theta and alpha bands, the power did not differ between the rTMS and sham conditions. In beta and gamma bands, the difference was marginally significant, F(1, 13) = 4.32, p = 0.058, and F(1, 13) = 3.56, p = 0.082, for the main effect of Stimulation in the beta and gamma bands, respectively.

**Figure 3.**
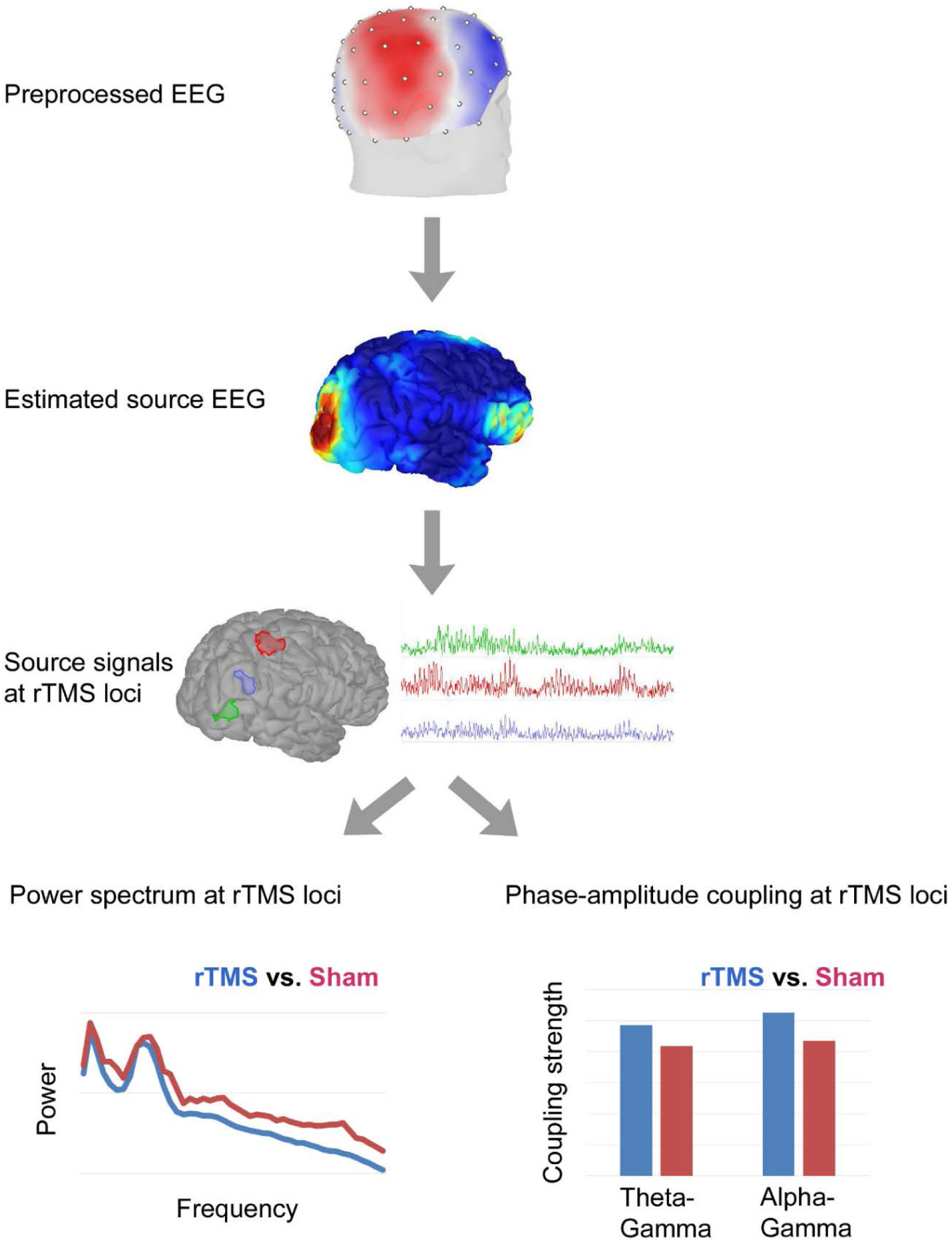
Illustration of the data analysis flow. Preprocessed EEG sensor-level signal was converted to source level signal. Signals from three sTMS loci were used for spectrum and phase-amplitude coupling analyses.

**Figure 4.**
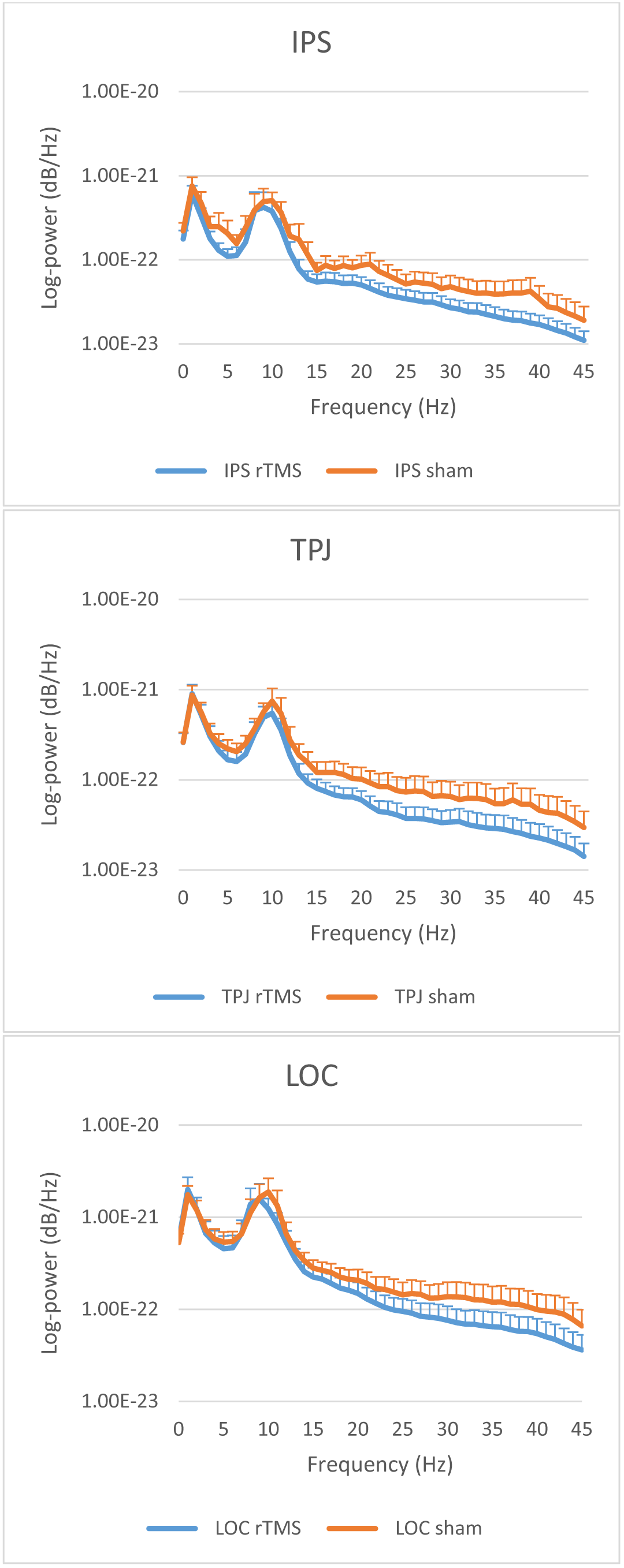
Power spectra in rTMS and sham condition in IPS, TPJ and LOC regions. Line graphs indicate grand average of power spectrum in rTMS (blue) and sham (orange) conditions at right IPS, TPJ, and LOC. Y axis is in log scale. Error bars indicate SE.

Since the rTMS frequency was 10 Hz, this could modulate the frequency of intrinsic alpha activity. To test if such effect had occurred, peak alpha frequency was taken from each participant. The maximum power value between 7 and 15 Hz was considered as the alpha peak frequency. In the IPS region, the peak value was, on average, 9.79 and 9.86 Hz for the rTMS and sham conditions, respectively; in the TPJ region, the average was 9.93 and 9.71 Hz for the rTMS and sham conditions, respectively; and in the LOC region, the average was 9.93 and 9.79 Hz, respectively. A 2-way (Stimulation by Loci) ANOVA was performed to test the differences among the averages. The results showed that no difference was statistically significant, all Fs < 1.

In sum, these analyses showed that the effect of rTMS on the power spectrum of the signal was restricted to a decreasing tendency in the fast bands.

### Phase-amplitude coupling (PAC) analysis

#### Selection of phase and amplitude frequencies

In the power spectrum of every participant, a peak was observed between 8 and 12 Hz. Thus, the signal between 8 and 12 Hz, i.e., alpha band, was used as the primary phase-providing band. In the theta band, a peak was visible in some but not all participants. Based on the observed peak locations and the PAC literature, in which the theta band has been discussed as the major phase-providing band [18, 29, 45], signals between 5 and 7 Hz were used to compute theta phase.

The amplitude band of interest was the gamma band (> 30Hz). To avoid muscular and mains artefacts, the bandwidth was limited between 30 and 45 Hz. In principle, an intermodulation component of gamma and alpha/theta activity could exist outside of this range, e.g., 20Hz, which is the f_2_-f_1_ component of f_1_ = 10 Hz and f_2_ = 30 Hz. To include such components in PAC estimation, a wideband signal, e.g., 15 - 45 Hz, could be chosen. In this case, however, the lower part of the wideband signal overlaps with the beta band activity. Moreover, the first harmonic of the alpha band activity peaks around 20 Hz. These non-gamma/non modulatory components in the amplitude signal could impair PAC estimation. Our choice to use the signal between 30 and 45 Hz for computing the amplitude signal circumvents these problems.

The phase and amplitude bands are sufficiently separated in frequency for PAC estimation. Central frequencies of the phase, f_1_, and amplitude bands, f_2_, satisfy; f_2_ / f_1_ > 2, as recommended [46].

#### Estimation method

PAC was estimated using Canolty’s method [29]. The original MATLAB code was adapted to Python (ver. 2.7). The single epoch signal was band-passed for 5 - 7 Hz, 8 – 12 Hz and 30 - 45 Hz for theta, alpha and gamma bands, respectively. Hilbert transform was applied to the band signals to compute instantaneous phase and amplitude. The phase and the amplitude signals were combined as:

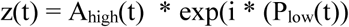

where P_low_(t) is the phase and A_high_(t) is the amplitude. The complex time series z(t) was averaged (MEAN_raw_) over a single epoch. The epoch contain approx. 36 cycles of theta activity (∼6 Hz, the slowest phase activity). In a previous study, we reported that the variance of PAC estimation decreased as the number of phase cycles in estimation increased, and became stable after about 10 phase cycles [32]. The current epoch length was, therefore, sufficient. Non-zero MEAN_raw_ indicates dependence between the phase and amplitude. For statistical evaluation, from each segment, 200 surrogate series were obtained using Canolty’s method. The MEAN_raw_ was normalized as:

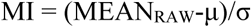

where μ and σ were the mean and variance, respectively of MEAN_surrogate_. Thus, MI is a z value (so the unit of MI is arbitrary) to be used for deciding on the presence of PAC. We used the 95 percentile of the z value, 1.65, as the reference value for the coupling.

#### TMS-evoked and noise-evoked activity

Our pilot study showed that single-pulse TMS evoked activity returned to baseline before 500 ms. Similarly, evoked activity due to the clicking noise in the sham condition returned to baseline before 500 ms. The evoked activity was avoided since data segments were analyzed between 500 and 9500 ms from the last pulse of rTMS/click noise.

### PAC results

Figure 5 shows the group mean of the medians in each stimulation site. The mean MIs were higher than the reference value for PAC, z = 1.65 (p < 0.05) in all conditions. Moreover, the mean MI appear to be higher in the rTMS than the sham conditions. A linear model was fit to the group data. The model was three-way repeated measures; Stimulation (rTMS vs. sham) by Loci (IPS, TPJ, and LOC) by Phase band (theta and alpha). The main effect of Stimulation was significant, F(1, 13) = 48.84, p < 0.00001. The main effects of Loci, Phase bands, and interactions were not significant, Fs < 1.1.

**Figure 5.**
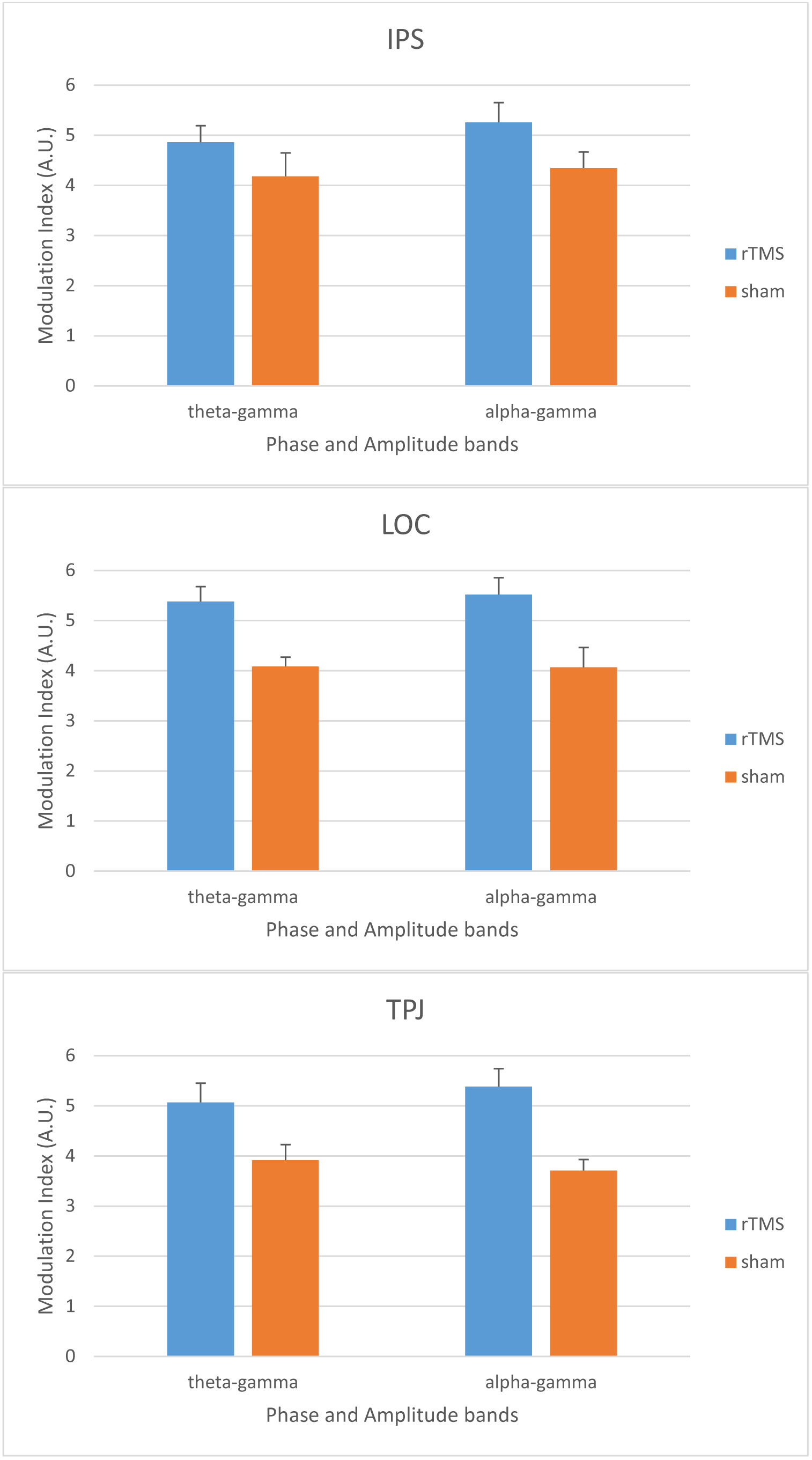
Phase-amplitude modulation index (MI) in group average. Bar graphs indicate grand average of MI in rTMS (blue) and sham (orange) conditions at right IPS, TPJ, and LOC. Band pairs are indicated in the horizontal axis. Error bars indicate SE.

The MI was compared between the first and the last epochs to check the effect of repeated application of rTMS. Since some epochs were omitted due to noise, the first epoch was the 1^st^ or the 2^nd^, and the last trial was one between the 12^th^ and 20^th^ of the actual 20 rTMS epochs. A four-way repeated measures model, Order (first vs. last) by Stimulation by Loci by Phase bands, was fit to the data. The main effect of Order was not significant, F<1. The main effect of Stimulation was significant, F(1, 13) = 8.54, p = 0.012, however, its interaction with Order was not, F<1. Another two-way interaction, Order by Phase bands, F(1, 13) = 4.52, p = 0.053, and a three-way interaction, Order by Phase bands by Loci, F(2, 26) = 3.82, p = 0.035. Between the 1^st^ and the last epochs, theta-gamma PAC decreased, while alpha-gamma PAC increased in IPS and LOC. In TPJ, however, both PAC decreased (Figure 6).

**Figure 6.**
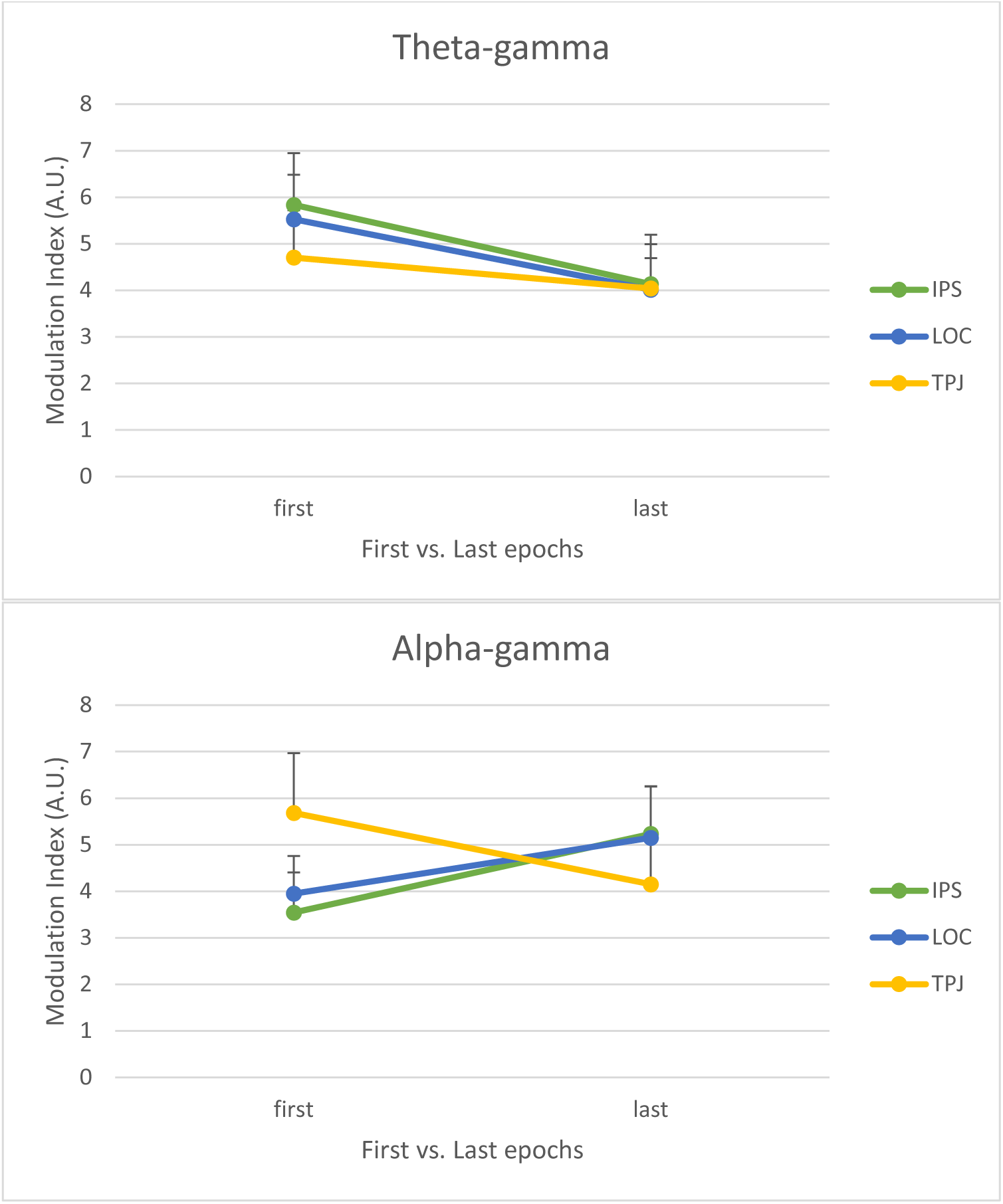
PAC in the first vs. the last epochs. Group mean and SE are plot for IPS (green), LOC (dark blue), and TPJ (yellow)

### Phase synchrony analysis of the phase-providing bands

In principle, the two phase-providing bands, theta and alpha, could also modulate each other. For example, [18] proposed hierarchical control from slower to faster activity, e.g., theta modulates alpha. The two phase-providing bands in the current study were too close in frequency to estimate their modulation using PAC (central frequency of theta, f_1_ = 6 Hz and central frequency of alpha f_2_ = 10Hz does not satisfy f_2_ / f_1_ > 2, in Aru et al., 2014). Thus, instead of PAC, cross-frequency phase-phase synchrony between the bands was estimated. The analysis was implemented in Python. First, a cycle of theta activity was binned to 16 bins. Alpha phase was sorted by the theta phase bins. In each bin, more than 250 alpha phase values were obtained. The circular mean of the alpha phase values was computed in each bin. Bin means (vector lengths) were used to plot a phase-phase histogram. Second, the deviation of the histogram from the uniform distribution was checked. If the histogram deviates from the uniform distribution, the phase-phase coupling between theta and alpha is not random. The deviation was estimated by Kullback–Leibler divergence (KLD). The KLD between the phase-phase and Uniform distribution was defined as:

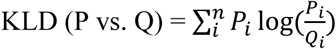

Where P is the normalized phase-phase histogram, Q is the histogram of a uniform distribution, and i is the number of bins. To obtain a probability distribution for the observed KLD, two hundred surrogate data sets were generated by splitting the alpha phase signals at a time point and swapping the split intervals. The splitting point was chosen randomly between 150ms and the end of a segment. The 200 surrogate KLD values were used as the bootstrap probability distribution for the original (observed) KLD. The 95 percentile value of the surrogate KLDs was taken as the threshold to judge non-randomness of the phase-phase coupling. The surrogate analysis was performed in single epochs. Less than 1% of all epochs showed non-random phase-phase coupling. It follows that phase synchrony between the phase-providing bands was negligible in the current data.

## Discussion

We investigated the effect of rTMS on theta/alpha-gamma PAC. Effects of rTMS were hypothesized based on a neural mass model (Chehelcheraghi, et al., 2016). The model has two control parameters for PAC generation; One is input noise level, which determines the emergence and stability of gamma oscillation, and the other is connectivity between pyramidal/slow inhibitory interneuron and fast inhibitory interneuron populations, which determines the strength of modulation. These parameters jointly determine PAC intensity observed in field signals. Of these, the connectivity is the primary parameter for persistent modulation on PAC, which is our current focus. The rTMS was applied to change the connectivity via Hebb’s principle; repeated and forced simultaneous firing increases the connectivity. As the connectivity increases, PAC should also increase. Results showed that the PAC between theta/alpha and gamma band activities increased in the post-rTMS period (∼10s), in accordance with the model prediction.

PAC increased in all stimulated regions. The regions, right IPS, TPJ, and LOC, were chosen as stimulation loci based on our interest in high-level mental functions. IPS has been regarded as the key region for multi-modal information integration for representation of external world for action [47]. TPJ has been associated to theory of mind and to integration of global and location information in visual scenes [48]. LOC has been considered critical for visual object recognition [49-51]. In spite of differences in functional implications, all regions responded similarly to rTMS in terms of PAC. This suggests that the neural mechanisms for PAC are similar across regions. Consequently, for modeling of PAC phenomena, the same model architecture could be applied across regions (fine-tuning of parameters could be needed for individual regions, e.g., [52]).

By definition, the high-level processes in these regions integrate information from low(er)-level processes. Therefore, modulation should sustain long enough to affect to the high-level functions. Our results show that the rTMS method can realize sustained PAC modulation, at least for 10s or so. Ten seconds which the current method can modulate PAC is long enough for a high-level function to be deployed in these regions. Also, it is long enough for performing a high-level cognitive task (e.g., Attentional Blink task in Nakatani et al., 2014). The current rTMS method, therefore, can be applied to experimentally investigate the relationship between PAC and cognition. How much further the effect could persist is currently unknown. Previous studies have been reporting persisting rTMS effects on various EEG and behavioral measures, of which the duration is often between several to 30 minutes (e.g., [40, 53]).

As the number of the stimulations increases, PAC strength would increase until the connectivity reaches to the maximum. In terms of accumulation of rTMS effect, subtle differences were observed among the regions and modulating (phase-providing) activities; whereas PAC increased in all loci and bands relative to the sham condition, as rTMS was repeatedly applied, alpha-gamma PAC increased relative to PAC of the first post-rTMS epoch, but theta-gamma PAC decreased in IPS and LOC. In TPJ, both decreased. The results suggest that an asymptotic function for connectivity could explain only part of the results. Factors such as neural fatigue, synaptic plasticity dynamics [54], and regional dominance of phase-providing bands [3] may also play a role in the long-term effect of rTMS on PAC.

Between the two phase-providing bands, a hierarchical relation of control has been suggested [18]. Our phases-phase synchrony analysis, however, did not reveal any specific relationship. [45] reported that the dominant modulator/phase-providing band shifted depending on task. In their study, PAC was computed between theta/alpha and high-gamma (> 50 Hz) bands, using ECoG from epilepsy patients who underwent a pre-surgical monitoring period. When the patients performed a visual task, PAC was stronger in alpha-high gamma than in theta-high gamma pairs. During a non-visual task, however, no significant difference was observed between the phase bands. Moreover, this task-specific effect was observed in lateral-and medial-temporal regions, but not in frontal regions. In our current study, the only effect of phase-providing bands (the phase by epochs interaction) was also region-specific. Taken together, these results suggest that gamma activity is modulated by theta and alpha bands which are uncorrelated to each other, and their modulation strength could change depending on tasks and brain regions.

## Conclusions

To increase PAC persistently in cortical regions, rTMS is shown to be effective. The observed effect of rTMS on the PAC suggests that spontaneous slow activity increased the coupling with fast activity. According to the model, the effect is due to the increased connectivity between the pyramidal and fast-responding interneurons. The effect was observed in right IPS, TPJ, and LOC, which suggests that similar mechanism operate in these posterior cortical regions for the modulation of the gamma activity reflecting the collective spiking behavior of pyramidal neurons [7].

## Acknowledgements

We would like to thank to Profs. Patrick Dupont and Rik Vandenberghe of the Laboratory for Cognitive Neurology at the University Hospital Gasthuisberg, who generously allowed us to use their experimental facility. We would also like to thank Dr. Maarten Schrooten of the Research Group Experimental Neurology at the hospital and Ms. Nicky Daniels of the Brain and Cognition Research Unit at KU Leuven for their technical support.

The study was jointly supported by research grants from the Flemish Government awarded to Johan Wagemans (METH/14/02) and to Cees van Leeuwen (Odysseus).

